# TaxonGPT: Taxonomic Classification Using Generative Artificial Intelligence

**DOI:** 10.1101/2024.10.28.618575

**Authors:** Haoyuan Huang, Teng Li, Zhixuan Wang, David Seldon, Allen Rodrigo

## Abstract

To address the challenges of consistency and quality in taxonomic classifications, we have developed a Python program called TaxonGPT, that utilizes the natural language processing capabilities of generative artificial intelligence (gAI, specifically, ChatGPT-4o) to generate taxonomic descriptions and taxonomic keys. To counter the propensity for large language model gAIs to “hallucinate”, we use knowledge graph semantic representation and an error-checking module to ensure that accurate taxonomic descriptions and keys are obtained. In this paper, we describe how TaxonGPT embeds ChatGPT-4o’s responses as outputs. We also report on benchmark tests for accuracy, efficiency and reproducibility. These tests demonstrate that TaxonGPT excels in generating taxonomic keys and descriptions.

## Introduction

Although biological taxonomy has well-established workflows, it is estimated that only 10% of species have been fully named or described taxonomically (Mora et al., 2011). Bacher (2012) pointed out that despite the exponential increase in the number of taxonomists, these numbers cannot keep up with the uptick in species extinction rates, resulting in many species potentially going extinct before being described. Thus, current taxonomic workflows and the number of taxonomists is insufficient to meet present taxonomic demands (Agnarsson and Kuntner, 2007; Pearson et al., 2011; Costello et al., 2013). In 2009, La Salle et al. (2009) discussed what they referred to as the ”Taxonomic impediment” at the Convention on Biological Diversity (CBD), highlighting three main dilemmas: uneven distribution of global taxonomic resources, lack of unified data standards, and over-reliance on taxonomists’ expertise (Rodman and Cody, 2003; Dar et al., 2012). Specifically, taxonomists in some regions lack sufficient resources to conduct research, and the absence of consistent data standards hinders the aggregation and sharing of taxonomic data.

Integrating information technology with taxonomy may provide an effective solution to the challenges of inconsistent and sometimes arbitrary choices made in recording data, and in describing and diagnosing taxa. This is not a new idea: for some time, information technology has been pivotal in managing taxonomic data, with international public domain databases marking the advent of data sharing (Green, 1994; Greenberg et al., 2009). The establishment of taxonomic research databases to collect and share data alleviates the uneven distribution of resources, often standardizes data formats, and facilitates better exploration of evolutionary relationships among species (Page, 2005; Miralles et al., 2020).

In 2000, DELTA (DEscription Language for TAxonomy) was developed as an information language system specifically designed for biological taxonomy (Coleman et al., 2010). The DELTA software provides a format for encoding taxonomic descriptions in a structured manner to record species characteristics. The system also contains extensive taxonomic data in a standardized DELTA format, assisting users in describing, storing, retrieving, and analyzing taxonomic information on biological species. With its standardized data format, DELTA can generate taxonomic keys, descriptions, and other outputs, making it a crucial tool for recording and analyzing data in taxonomic research (Whitfield et al., 2009; Sharkey et al., 2009, 2011; Esmaeili-Rineh et al., 2015; Fišer et al., 2018).

Over the last few years, a revolution in generative artificial intelligence (gAI) and the use of language models trained on very large corpuses of text has ignited significant interest in gAI’s utility across a range of scientific areas and disciplines (Liu et al., 2023; Gangwal et al., 2024). A Large Language Model (LLM) is a highly parameterized neural relevant fragments of words and text) in a sentence, and evaluates the co-occurrence amongst tokens by taking account of context (Li et al., 2022; Katz et al., 2022; Birhane et al., 2023). These models facilitate gAI’s general language capabilities by applying deep learning techniques to learn structural language patterns and semantic relationships from extensive text data. Large-scale neural language models, which are pre-trained on extensive text data and then fine-tuned for specific tasks, have achieved significant success in various general-purpose natural language processing (NLP) tasks (Beltagy et al., 2022; Tan et al., 2023; Zhang et al., 2024). Thus, LLMs are able to produce apparently meaningful, human-comprehensible textual responses to queries written in natural language, and are now widely used in tasks such as text generation, automatic translation, and speech recognition (Brown et al., 2020; Gu et al., 2022).

In biology, Gu et al. (2022) have demonstrated that pre-trained large language models also excel in specialized domains, achieving or even exceeding human-level performance in reading comprehension and question-and-answer tasks in biomedicine, accurately answering specialized biomedical knowledge (Beltagy et al., 2019).

ChatGPT-4o, the latest Generative Pre-Trained Transformer (GPT) model developed by OpenAI (San Francisco, California, USA; http://openai.com), can extract meaning from complex queries that may be constructed in different ways in a variety of natural languages, and is able to respond appropriately (Shahriar et al., 2024; Sonoda et al., 2024). Since taxonomy relies largely on the use of natural language to describe taxa (Gregg, 1950), and the attendant character states that each taxon possesses, it is reasonable to suggest that LLMs and GPTs like ChatGPT-4o offer ways to improve both the consistency of taxonomic data handling, and the descriptive and diagnostic tools to address the taxonomic impediment.

It is important to note that although GPT language models have demonstrated excellent results in NLP tasks and may have significant potential in taxonomy research, they also are known to produce ”Artificial Intelligence (AI) hallucinations” (Feldman et al., 2023), i.e., responses that are irrelevant, factually incorrect, illogical, or contradictory (Perković et al., 2024). This presents an obvious challenge in any AI application to taxonomy since it is imperative that taxonomic descriptions faithfully describe the taxa in question. This extends to ensuring that character data are collated uniformly, and reported accurately and consistently.

To this end, AI researchers have demonstrated that the use of data structures that explicitly define relationships between semantic elements go a long way towards minimizing AI hallucinations (Brown et al., 2020; Rae et al., 2021). A knowledge graph——an approach that we apply here——is one such structure. A knowledge graph is a structured semantic graphical knowledge base that provides additional information about the relationships between elements (in our case, between taxa and character states), so that GPT language models understand data more accurately and avoid factual errors by referencing structured data patterns (Mohamed et al., 2020).

In this paper, we apply OpenAI’s GPT (Generative Pre-Trained Transformer) language model (ChatGPT-4o) to two taxonomic tasks: the generation of taxonomic descriptions and the creation of taxonomic keys, based on the morphological character matrix with corresponding morphological character information.

Specifically, TaxonGPT is a Python program that utilizes the most recent version of ChatGPT at the time of writing (Fig. 1), ChatGPT-4o (06/08/2024), to produce taxonomic descriptions and taxonomic keys. It does this by making calls to ChatGPT-4o’s Application Programming Interface (API), and extracting appropriate natural language responses. TaxonGPT consists of three modules: (a) DESCRIBE, a module for generating taxonomic descriptions using the Nexus matrix file with corresponding morphological character information; (b) KEY, a module for generating taxonomic keys using the Nexus matrix file, and (c) CORRECT, an error-correcting module which is designed for the KEY module. CORRECT uses the self-learning error correction capabilities of ChatGPT-4o, by feeding back erroneous results into the GPT for correction. This iterative process ensures the accuracy of the final outputs. With ChatGPT-4o, we demonstrate that TaxonGPT can provide high-quality descriptions and diagnoses that are consistent and accurate. To run TaxonGPT, users need to download and install **TaxonGPT.py**, a Python program, on a local machine. TaxonGPT requires an Application Programming Interface (API) Key from OpenAI, and this is available with a charge.

**Fig. 1.**
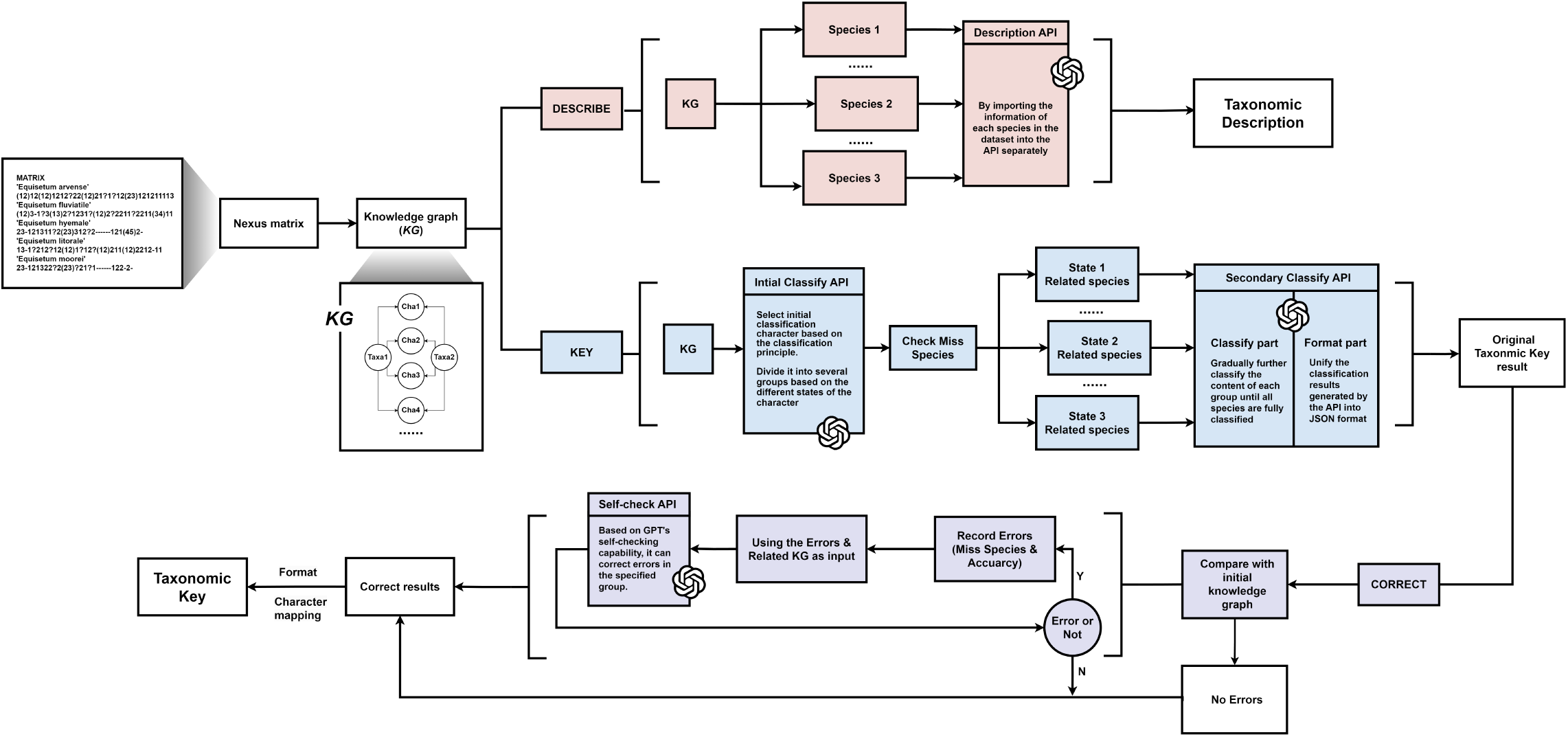
Flowchart showing the functional modules in the TaxonGPT: Use OpenAi company annotations in every place where the API model is called.

## Description

### Architecture of TaxonGPT

TaxonGPT has three main modules: DESCRIBE, KEY, and CORRECT (Fig. 1).

The user can run DESCRIBE and KEY independently, by providing a common configuration (config) file. The config file specifies path names for the Nexus morphological matrix, corresponding morphological character information (i.e., character names and states) which can be integrated within the morphological matrix or provided separately, a pathname for a prompt file in editable Javascript Object Notation (JSON) format, an OpenAI API key, and output file pathnames for a CSV formatted morphological matrix, knowledge graph-formatted morphological matrix, a taxonomic description file and taxonomic key file. CORRECT is an error-checking module that runs with KEY and is not available to the user as an independent module.

The flowchart in Figure 1 describes the operation of modules DESCRIBE, KEY, and CORRECT. A pre-processing step converts the Nexus matrix and the character state files into a Knowledge Graph (KG). *KG* is then used as input for each module. DESCRIBE (in Red) generates taxonomic descriptions by calling the API of ChatGPT-4o several times, and utilizes *KG* to generate complete descriptions. KEY (in Blue) generates taxonomic keys by using multiple API calls, to iteratively refine the key. After a full taxonomic key is produced, errors are recorded by comparing *KG* to the key using CORRECT. CORRECT reports errors back to ChatGPT-4o, resulting in error correction at the appropriate iteration round used by KEY. To communicate with ChatGPT-4o, TaxonGPT uses natural language prompts stored in a JSON file. By organizing the prompts within a hierarchical structure, API calls using appropriate prompts can be made at relevant points in the execution of TaxonGPT. Since the JSON file is text-readable, researchers can edit, customize, and configure the prompts to meet their specific research requirements. Detailed descriptions of DESCRIBE, KEY, and CORRECT can be found in Supplementary Materials 1.

### Evaluation of TaxonGPT

#### Trial Datasets

Datasets from DELTA (Descriptive Language for Taxonomy) (Askevold and O’Brien, 1994) were used to evaluate TaxonGPT. Based on extensive testing, we noted that ChatGPT-4o model is unable to analyze datasets with more than 30 species. Due to this limitation, six datasets were finally selected from the DELTA database, and five additional taxonomic datasets were obtained from published literature (Whitfield et al., 2009; Sharkey et al., 2009, 2011; Esmaeili-Rineh et al., 2015; Fišer et al., 2018), for a total of 11 taxonomic datasets that included Nexus morphological matrices and morphological information, for evaluation. Additionally, because ChatGPT-4o can provide alternative responses to queries, we assessed TaxonGPT by evaluating five responses for each dataset to determine the accuracy and consistency of the pipeline.

#### Comparison to other software

The results generated by TaxonGPT were compared with those produced by the web version of ChatGPT-4o and DELTA. This comparison was conducted to assess TaxonGPT’s performance in terms of classification quality, accuracy, efficiency, and reproducibility. The web version of GPT-4o refers to the utilization of the ChatGPT-4o via the ChatGPT web page (https://chatgpt.com), wherein relevant prompts (Supplementary Materials 2) and raw taxonomic data are submitted online to generate the corresponding classification results.

#### Accuracy Measures

The accuracy of the taxonomic results (taxonomic descriptions and taxonomic keys) generated by TaxonGPT was assessed by comparing the morphological characters of species in the original data with the final classification results, with a focus on identifying any errors in morphological character states. The error rate was determined by calculating the ratio between correctly and incorrectly used morphological characters in the taxonomic results, and an average was taken from the five repeated trials. This error rate serves to quantify the accuracy of the final taxonomic results, thereby providing an evaluation of the reliability and accuracy of the results generated by TaxonGPT.

#### Taxonomic Key Quality

The *E*_Dicho_ index was utilized to quantify the quality of the taxonomic key results (Sinh et al., 2017). The *E*_Dicho_ based on Pielou’s Evenness Index (Pielou, 1966), was used to evaluate the evenness and classification effeciency of dichotomous taxonomic keys. Higher *E*_Dicho_ values indicate more efficient taxonomic keys, i.e., keys with fewer average steps to classifying each taxon. The *E*_Dicho_ index is calculated using the formula:

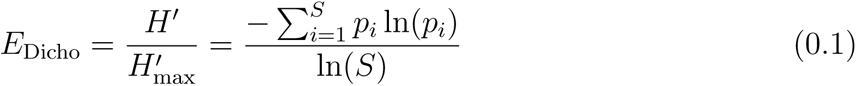

Where *H*′ is the Shannon diversity index calculated as 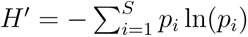, where *p_i_* represents the proportion of steps required to identify each taxon, and *S* is the total number of taxa in the key. This method is suitable for evaluating both dichotomous and general taxonomic keys. The *E*_Dicho_ values ranges from 0 to 1, where *E*_Dicho_ = 1 indicates that the taxonomic key provides the most efficient and balanced discrimination among all taxa on average. As noted above, to assess TaxonGPT, taxonomic keys were generated for 11 datasets, with five repeated trials conducted for each dataset. Subsequently, the *E*_Dicho_ index was calculated for each of the five repeated results of each dataset and an average was determined.

In the Nexus matrix, some species’ morphological character states may be marked as ”Missing” or ”Not applicable”(Pleijel, 1995; Maddison et al., 1997). To further assess the quality of the taxonomic key generated by TaxonGPT, the presence of these special character states (marked as ‘Missing’ or ‘Not applicable’) in the taxonomic keys was evaluated. ”Missing” denotes that the character state information for the current taxon is unavailable, possibly due to data loss or incomplete data collection. Such data is deemed invalid because it provides no useful taxonomic information. In contrast, ”Not applicable” pertains to a character state that is irrelevant or inapplicable to a particular taxon but still conveys meaningful information. For example, consider the character ”number of legs” when constructing a key that includes both vertebrates and invertebrates. Invertebrates like octopuses have eight arms, not legs, making the character ”number of legs” inapplicable. In this case, ”Not applicable” is the appropriate state because it accurately reflects the biological reality of the taxon in question. This information is valid and useful, as it helps differentiate octopuses from other animals in the taxonomic key (Wilkinson, 1995; Strong and Lipscomb, 1999). Thus, ”Missing” is regarded as an invalid state, whereas ”Not applicable” is recognized as a valid state. In the 11 datasets we have used to evaluate TaxonGPT, “Missing” is denoted by “?” and “Not applicable” is denoted by “—”. By tallying the frequency of invalid character states used in the taxonomic keys and comparing it to the total frequency of morphological character states used, the quality of the taxonomic key can be assessed.

#### Reproducibility

To assess the reproducibility of the results generated by TaxonGPT, five repeated trials were conducted for each of the 11 datasets, producing classification outputs (taxonomic descriptions and taxonomic keys). To evaluate the consistency of the morphological characters extracted in the taxonomic keys across multiple repeated experiments, a Consistency Score (CS) was calculated. This score is based on the number of shared features between the feature list from the output of the first trial, and each of the feature lists from the subsequent four trials. We chose to compare all runs to the first, because users would typically rely on the output of the first iteration, when reporting results.

Let the baseline feature list be *C*_1_, and the comparison feature lists be *C*_1_ (where *i* =2,3,4,5). Thus, the Consistency score *CS* for each dataset is the average of these four scores:

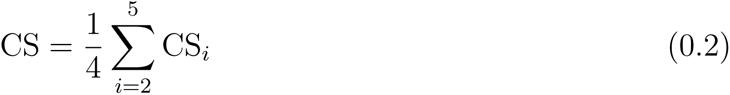

Where *CS_i_*is calculated as:

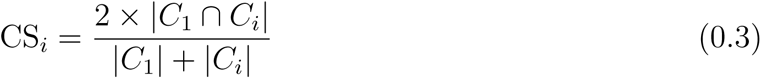

And:

*• |C*_1_ *∩ C_i_|* represents the number of shared features between the baseline feature list *C*_1_ and the comparison feature list *C_i_*.
*• |C*_1_| is the total number of features in the baseline feature list *C*_1_.
*• |C_i_|* is the total number of features in the comparison feature list *C_i_*.

The consistency score *CS* ranges from 0 to 1, where *CS* = 1 means that the character lists are identical between the baseline and all other replicates. As a rough guide, *CS ×* 100 approximates the average percentage of character states shared with the baseline across all other iterations.

#### Efficiency and cost

The computational efficiency of TaxonGPT in producing classifications and the impact of the numbers of species and characters in the Nexus matrix on runtime was assessed by recording the time taken for each invocation of the ChatGPT-4o model to produce taxonomic descriptions and taxonomic keys. For all datasets, five repeated trials were performed.

To evaluate the costs incurred in generating different classification results (taxonomic description and taxonomic key) by TaxonGPT, as well as the impact of the number of species and morphological characters on the cost of generating classification results via the API, the costs of each invocation of the ChatGPT-4o model API to generate different classification results were recorded separately. As before, for all datasets, different classification results were generated independently, and again, five repeated trials were conducted, with the costs recorded each time. These data allow for the analysis and evaluation of the cost-effectiveness of TaxonGPT under various conditions.

## Benchmark

This section presents a comparative analysis of the classification results generated by TaxonGPT (API) and the web-based GPT-4o (Web) against those produced by existed taxonomic software (DELTA).

### Taxonomic Key Quality and Accuracy

The quality of the taxonomic keys was assessed using the *E*_Dicho_ index. The results indicated that the *E*_Dicho_ indices for TaxonGPT (API) and web-based GPT-4o (Web) (Fig. 2A) were generally higher than those for the DELTA software, suggesting that the ChatGPT-4o model is superior at generating high-quality taxonomic keys. Specifically, the web-based GPT-4o (Web) was found to achieve the highest average *E*_Dicho_ scores across all datasets: Web (0.990) *>*API (0.982) *>*DELTA (0.968).

**Fig. 2.**
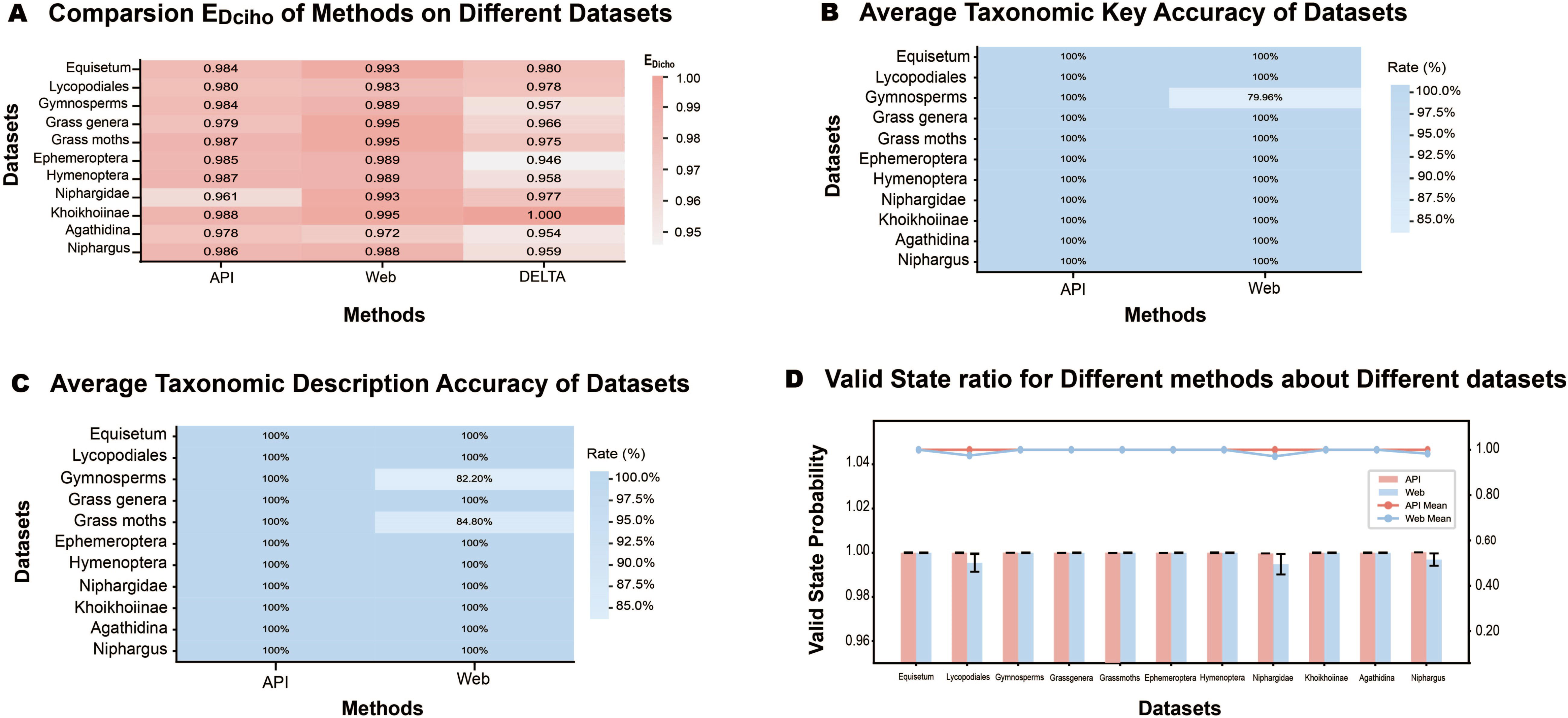
Benchmarks Performance Evaluation (Quality and Accuracy). **A.** Mean *E*_Dicho_ Scores: Comparison of average *E*_Dicho_ scores for classification results from API (0.982), Web (0.990), and DELTA (0.968) across all datasets. **B.** Taxonomy Key Accuracy: Comparison of average accuracy of Taxonomic key generated by API and Web. **C.** Taxonomic Descriptions Accuracy: Comparison of average accuracy of taxonomic descriptions produced by API and Web. **D.** Valid Classification States Probability: Comparison of the probability of valid classification states in results generated by API and Web across various datasets.

The presence of invalid character states (‘Missing’ state) in the taxonomic keys (Fig. 2D) was also assessed. A comprehensive assessment and comparison revealed no missing species in any of the generated classification results. The frequency of invalid states in the taxonomic keys was found to be relatively low (99.8%) when using the web-based GPT-4o (Web). In all 11 datasets, the proportion of valid states for TaxonGPT (API) was 100.0%. Invalid character states in the classification results from ChatGPT-4o (the web-based GPT-4o) were typically observed when analyzing datasets containing a larger number of characters (i.e., when the number of characters was greater than 30).

In assessing the accuracy of the generated taxonomic results (Fig. 2B), classification results were compared with the character states in the original taxonomic data, and the frequency of errors in the selected diagnostic character states was calculated. The taxonomic key produced by TaxonGPT was always correct (i.e., 100% accurate); in contrast, the web-based GPT-4o had difficulty accurately generating taxonomic keys for one large Gymnosperm dataset (with 75 characters), achieving an average accuracy of only 79.9% across five repeated classification results.

TaxonGPT also demonstrated significantly higher accuracy in generating taxonomic descriptions compared to the web-based GPT-4o (Fig. 2C). An accuracy of 100.0% was achieved by TaxonGPT in generating descriptions. In contrast, the web-based GPT-4o demonstrated lower accuracy in some cases (average of 97.0%).

### Efficiency and Costs

Taxonomy typically involves extensive manual work, requiring significant time to retrieve and select appropriate characters to classify. TaxonGPT aids in the generation of descriptions and keys. For taxonomic keys, the results indicated that for the web-based GPT-4o (Web) method, the number of characters in the dataset was not significantly correlated with runtime. In contrast, for TaxonGPT (API), a correlation between the number of characters and the runtime was observed, with more characters resulting in longer processing times (Fig. 3A, 3D). Additionally, the number of species was found to significantly affect the time required for both TaxonGPT (API) and web-based GPT-4o (Web) methods to generate taxonomic keys, with the higher the number of species requiring longer runtimes. The average time to generate taxonomic keys was found to be 88 seconds for TaxonGPT (API) and 92 seconds for the web-based GPT-4o (Web) (Fig. 3C). Currently, GPT-4o is freely available to users in certain regions, allowing up to 30 messages to be sent to ChatGPT every three hours. For additional usage, a monthly fee of US$20 is required. In the Python package, the cost of using the GPT-4o model called by TaxonGPT was directly proportional to the number of characters and species being classified. In this study, the cost of generate each taxonomic key was found to be less than US$1 (Fig. 3G).

**Fig. 3.**
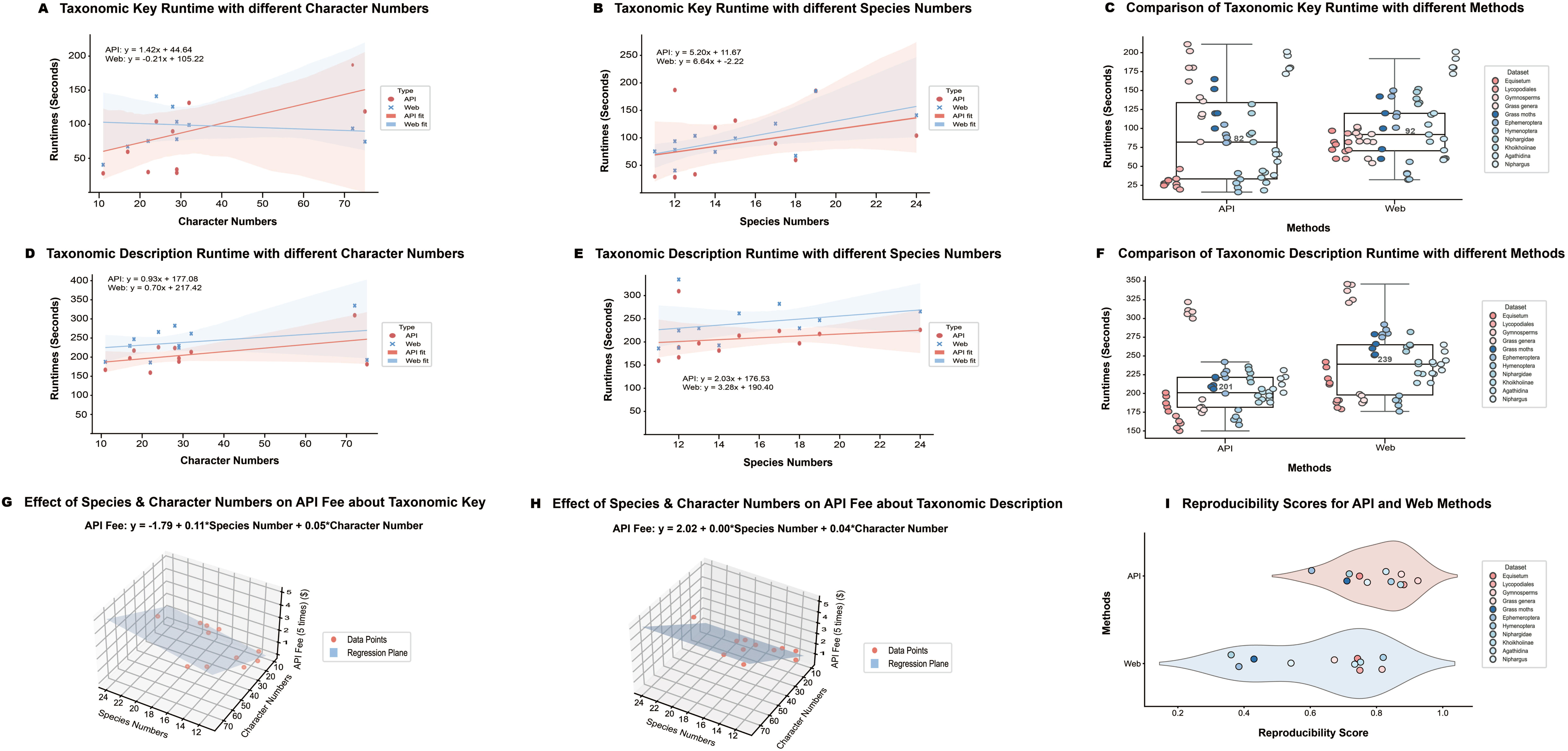
Benchmark Performance Evaluations (Efficiency, Costs and Reproducibility). **A.** Effects of different character numbers on the runtime of taxonomic keys produced by various methods (API and Web). **B.** Effects of different taxon numbers on the runtime of taxonomic keys produced by various methods (API and Web). **C.** Comparison of the runtime required to generate taxonomic keys using API and Web methods, with each box-plot representing the distribution of runtimes, including the median (central line), first and third quartiles (box edges), and data points within 1.5 times the interquartile range (whisker boundaries). **D.** Effects of different character numbers on the runtime for generating taxonomic descriptions produced by various methods (API and Web). **E.** Effects of different taxon numbers on the runtime for generating taxonomic descriptions produced by various methods (API and Web). **F.** Comparison of the runtime required to generate taxonomic descriptions using API and Web methods, regarding the specific explanation of the box plot, see above. **G/H.** Impact of the number of taxa and morphological characters on API costs for generating Taxonomic Key **(G)** and taxonomic descriptions **(H)**. **I.** Reproducibility Scores: Comparison of reproducibility scores for different datasets using API and Web methods.

For generating taxonomic descriptions, it is essential that comprehensive and biologically accurate descriptions are created, correctly linking species information to morphological characters. Consequently, both the web-based GPT-4o and TaxonGPT were impacted in runtime by the number of species and characters involved: the number of species and features were both positively correlated with the time taken to generate taxonomic descriptions (Fig. 3D, 3E). The average runtime for generating these descriptions was found to be 239 seconds for the web-based GPT-4o and 209 seconds for TaxonGPT (Fig. 3F). When generating taxonomic descriptions, the cost associated with TaxonGPT was higher compared to generating taxonomic keys due to the increased length of text generated. Nonetheless, the overall cost of generating taxonomic descriptions for all taxa within a dataset was found to not exceed US$1 per dataset (Fig. 3H).

### Reproducibility

Finally, multiple tests were conducted on different methods and datasets to assess the reproducibility of generating classification results with TaxonGPT (API) and the web-based GPT-4o (Web) (Fig. 3I). TaxonGPT (API) achieved a average consistency score, CS, of 0.8 in character state selection for the taxonomic keys, across the five replicates for each of the 11 datasets. The web-based GPT-4o (Web) achieved an average CS of only 0.65 across multiple tests.

## TaxonGPT Example Output

To illustrate the outputs of TaxonGPT, the dataset for the genus *Equisetum* was extracted from the DELTA database. This dataset comprises 29 morphological characters for 12 species, stored in the form of a Nexus matrix (Fig. 4).

**Fig. 4.**
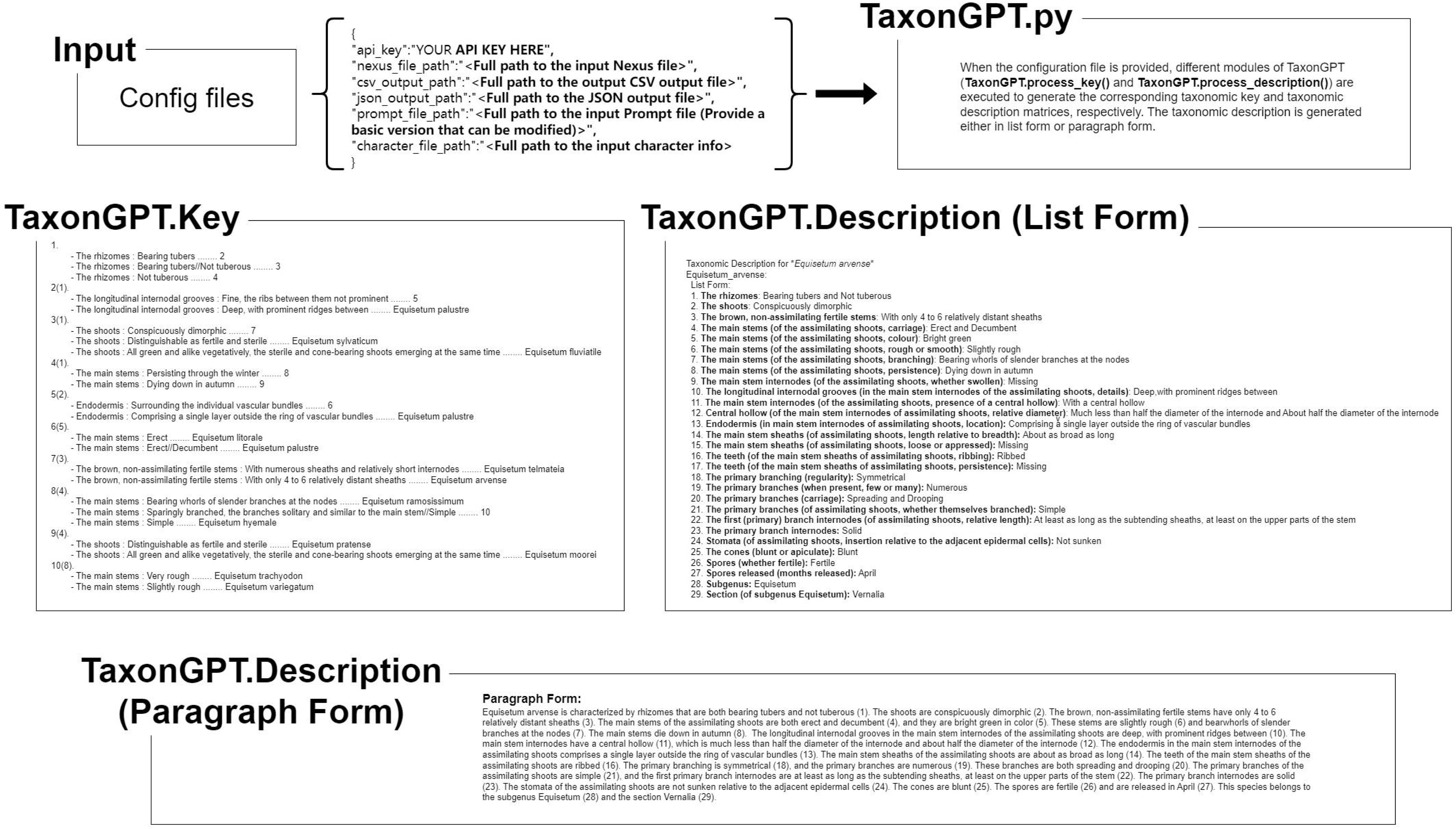
Examples of Taxonomic Results Generated by TaxonGPT. This figure illustrates a complete example of taxonomic results generated by TaxonGPT, along with the necessary inputs. The ”Input” section contains the configuration files required for TaxonGPT, including the API key for the ChatGPT-4o model, the relevant Nexus matrix file, and the corresponding morphological character information. TaxonGPT provides a base prompt file, which is modifiable by users in designated sections. After the inputs have been configured, executing TaxonGPT.process key () and TaxonGPT.process description () will generate the taxonomic outputs (a taxonomic key and a taxonomic description), with the latter available in both list and paragraph formats.

## Discussion

TaxonGPT is a Python program designed to parse Nexus morphological matrices and utilize the natural language processing (NLP) capabilities of generative AI, specifically ChatGPT-4o, to create taxonomic classifications. Through the flexible invocation of natural language output and the incorporation of knowledge graph data structures, TaxonGPT can generate consistent and accurate taxonomic descriptions and keys. When compared to traditional taxonomic software (e.g., DELTA), TaxonGPT offers a time-efficient, natural language-based and inexpensive alternative for one of the tasks that taxonomists perform.

However, there are certain limitations to the integration of large language models with taxonomy. TaxonGPT faces constraints in handling datasets with a large number of species and characters, primarily due to the computational limitations of current language models and the complexity of taxonomic relationships in large datasets. Although this issue can be mitigated by splitting large datasets and integrating partial classifications results, it remains challenging to process large-scale datasets without compromising accuracy or efficiency. We expect, however, that with the rapid improvements in gAI, this issue will be resolved soon.

While our assessments indicate that TaxonGPT provide high quality, consistent, and accurate descriptions and keys, these results still require professional review and appropriate editing before they can be applied directly. Taxonomists will still need to provide taxonomic names, where necessary, review information on authorship, check for synonyms, and so on. Therefore, TaxonGPT should be viewed as a tool to augment, rather than replace, expert knowledge in taxonomy.

As language models continue to evolve, natural language processing capabilities will improve. Future research should focus on training models on specialized taxonomic datasets to enhance their performance in this domain. Additionally, developing methods to handle larger datasets more efficiently and improving the interpretability of the AI’s decision-making process in classification tasks are crucial areas for advancement. As we have shown, generative AI has the ability to address the bottlenecks of the “taxonomic impediment” of La Salle et al. (2009). We are optimistic that the rapid improvements in generative AI will serve to accelerate systematics research, and provide the necessary tools to deliver consistent classifications.

## Acknowledgements

Special appreciation is extended to Andrea Grecu, June Ko, Yuan Xu, Jiancheng Li, Zhao Cao, and Miao Wang for their support and insightful discussions. For this manuscript, ChatGPT-4o assisted with language translation and with editorial assistance, including grammar and syntax corrections. The conception, methodology, analysis, and interpretation of results are exclusively the work of the authors.

## Supplementary Material

Data available from the Dryad Digital Repository: https://doi.org/10.5061/dryad.1vhhmgr3s.

Temporary sharing link: http://datadryad.org/stash/share/pwZ8nVTV3Gpt1fiUyUmjMUO9DTlOizDgl7rEQ3_2bPI.

## Data availability

The data used in this manuscript were sourced from publicly available repositories. Specifically, the trial dataset was obtained from the DELTA database and can be downloaded from the DELTA website (https://www.delta-intkey.com/www/data.htm). Additionally, taxonomic data stored in DELTA format from the following published studies were integrated into our analysis (Whitfield et al., 2009; Sharkey et al., 2009, 2011; Esmaeili-Rineh et al., 2015; Fišer et al., 2018). These data were primarily used for comparative analysis and verification of species character states. All data from these sources are publicly available through the respective publications. Any additional data generated during the course of this study are available upon request from the corresponding author.

## Code availability

TaxonGPT is provided as an open-source Python program, with a detailed user manual available in the Vignettes directory of the GitHub repository. All code used for TaxonGPT is publicly available, and the necessary usage files are also stored in a GitHub repository (https://github.com/hhua361/TaxonGPT-API). The analyses presented in this manuscript were performed using Python version 3.12.6.

